# In silico prediction of ARB resistance: A first step in creating personalized ARB therapy

**DOI:** 10.1101/2020.02.12.945402

**Authors:** Shane D. Anderson, Asna Tabassum, Jae Kyung Yeon, Garima Sharma, Priscilla Santos, Tik Hang Soong, Yin Win Thu, Isaac Nies, Tomomi Kurita, Andrew Chandler, Abdelaziz Alsamarah, Rhye-Samuel Kanassatega, Yun Luo, Wesley M. Botello-Smith, Bradley T. Andresen

## Abstract

Angiotensin II type 1 receptor (AT_1_R) blockers (ARBs) are among the most prescribed drugs. However, ARB effectiveness varies widely, and some patients do not respond to ARB therapy. One reason for the variability between patients is non-synonymous single nucleotide polymorphisms (nsSNPs) within *agtr1*, the AT_1_R gene. There are over 100 nsSNPs in the AT_1_R; therefore, this study embarked on determining which nsSNPs may abrogate the binding of selective ARBs. The crystal structure of olmesartan-bound human AT_1_R (PDB:4ZUD) served as a template to create an inactive empty AT_1_R via molecular dynamics simulation (n = 3). All simulations resulted in a smaller ligand-binding pocket than 4ZUD due to the absence of olmesartan in the simulation yet remained inactive with little movement in the receptor core. A single frame representing the average stable AT_1_R was used as a template to thrice (n = 3) dock each ARB via AutoDock to obtain a predicted affinity from the weighted average of 100 docking simulations. The results were far from known values; thus, an optimization protocol was initiated, resulting in the predicted binding affinities within experimentally determined ranges (n = 6). The empty model AT_1_R was altered and minimized in Molecular Operating Environment software to represent 103 of the known human AT_1_R polymorphisms. Each of the eight ARBs was then docked, using the optimized parameters, to each polymorphic AT_1_R (n = 6). Although each nsSNP has little effect on global AT_1_R structure, most nsSNPs drastically alter a sub-set of ARBs affinity to the AT_1_R. Comparisons to previous binding studies suggest that the results have a 60% chance of predicting ARB resistance. Although more biochemical studies and refinement of the model are required to increase the accuracy of the prediction of ARB resistance, personalized ARB therapy based on agtr1 sequence could increase overall ARB effectiveness.

**Author Summary:** The term “personalized medicine” was coined at the turn of the century, but most medicines are currently prescribed based on disease categories and occasionally racial demographics, but not personalized attributes. In cardiovascular medicine, the personalization of medication is minimal; however, it is accepted that not all patients respond equally to common cardiovascular medications. Here we chose one prominent cardiovascular drug target, the angiotensin receptor, and, using computer modeling, created preliminary models of over 100 known alterations to the angiotensin receptor to determine if the alterations changed the ability of clinically used drugs to interact with the angiotensin receptor. The strength of interaction was compared to the unaltered angiotensin receptor, generating a map predicting which alteration affected each drug. It is expected that in the future, a patient’s receptors can be sequenced, and maps, such as the one presented here, can be used to select the optimum medication based on the patient’s genetics. Such a process would allow for the personalization of current medication therapy.

## Introduction

The Angiotensin (Ang) II type 1 receptor (AT_1_R) is often studied due to its role in cardiovascular disease, diabetes, and, more recently, cancer.(1, 2) Eight clinically viable antagonists (ARBs) target the AT_1_R, and ARBs are widely used for the treatment of hypertension, heart failure, and chronic kidney disease. Additionally, due to the prominent role the AT_1_R plays in cardiovascular disease, the AT_1_R is also the focus of many genetic association studies.(3, 4) However, very few studies have been directed toward pharmacogenomics of the AT_1_R.

Not all patients respond equally, or at all, to ARBs.(5) One potential reason a patient does not respond, or respond optimally, to an ARB is that there could be a, or many, non-synonymous single nucleotide polymorphism(s) (nsSNP) within the *agtr1* coding sequence. nsSNPs can result in altered antagonist function;(6) thus, as we enter an era dominated by big data and likely personalized medicine, it would be ideal for a prescriber to know which therapies will interact with their target as expected in each patient. Such patient-specific knowledge can come from genetic screening coupled to robust databases linking drugs to effects. Alternatively, if there is no previous data, then there should be a mechanism allowing rapid assessment of which drugs are appropriate for a given patient.

The AT_1_R was cloned in the early 1990s and recently was crystallized with an ARB bound.(7, 8) Before crystallization, the ARB binding pocket was investigated primarily through mutagenesis studies.(9–12) These studies identified residues involved in ARB binding that are within the known binding pocket but also identified residues involved in ARB binding that are far from the known binding pocket.(9, 10) Such data demonstrate that single amino acid changes in the AT_1_R far from the binding pocket can alter the receptor conformation and disrupt ARB binding. Moreover, in many cases, the mutant AT_1_Rs still bound to, and transduced signals from, Ang II demonstrating the ability of an nsSNP to maintain the physiological functions of the AT_1_R yet display ARB resistance. Multiple genomic projects identified polymorphisms within the AT_1_R.(13) Herein, an empty AT_1_R was generated through molecular dynamic (MD) simulation, 103 nsSNPs from the 1000 genomes project were briefly modeled, and each of the eight clinically viable ARBs was docked to each AT_1_R in order to predict which nsSNPs would lead to ARB resistance. To our knowledge, this is the first large scale investigation into known human nsSNPs within the AT_1_R.

## Results

### Wild-type AT_1_R Model

Crystalized G protein-coupled receptors (GPCRs) often contain a tightly bound ligand that stabilizes a unique ligand-induced conformation. In order to obtain a model of the empty AT_1_R, the AT_1_R crystal structure (PDB: 4ZUD) was modified to remove non-receptor residues and the unresolved flexible loops were added back to the receptor, then a short 150 ns MD simulation was conducted within a POPC:cholesterol (87:13 ratio) membrane to relax the structure to a ligand-free state (n = 3). Each simulation reached a stable structure around 100 ns (Supplemental Figure 1A), and the last 20 ns of the simulations were stable. A single frame representing the average stable root-mean-square deviation (RMSD) from the last 20 ns of replica 1 was chosen as the representative model of the empty AT_1_R; the PDB file format is available in the supplemental files. A notable feature of the selected empty AT_1_R model is that helix 8 appeared to be an extension of helix 7 (Figure 1A). Therefore, the orientation of helix 8, in comparison to the model AT_1_R, was examined to determine its flexibility and spatial orientation (Supplemental Figure 1B). Throughout the simulations, the position of helix 8 varied by 12 angstroms from the empty AT_1_R model, which is similar to previous MD simulations.(14) To further address the overall movement of the simulations, distance measurements corresponding to double electron-electron resonance (DEER) spectroscopy of the AT_1_R(14) were measured over the last 20 ns of simulation (Figure 1B). Red Xs represent the major modes of the DEER data in Figure 1B; 50% of the major modes from the DEER experiments are within distances observed in the modeled empty AT_1_R. Moreover, most of the measurements from the model lie within the range identified in the DEER spectroscopy data set but capture a different state than the MD modeling presented with the DEER spectroscopy. Only F55-R139, which Wingler et al. call TM1-TM2, and D236-R311, called TM6-helix 8, do not necessarily represent the DEER spectroscopy. To capture the fluctuation within all residues, RMSF was plotted as the β-factor and colored based on the degree of movement (Figure 1A). The flexible loops, as well as helix 8, showed the most fluctuation; however, the cytosolic and extracellular helical interfaces also showed, on average, greater than 2 Å of movement. Expectedly, removal of olmesartan resulted in movement in the ligand-binding pocket (upper third of the AT_1_R); however, the remaining cores of the receptor displayed little movement. The data suggests that the empty AT_1_R model is a viable and unique model of the apo-AT_1_R.

**Figure 1:**
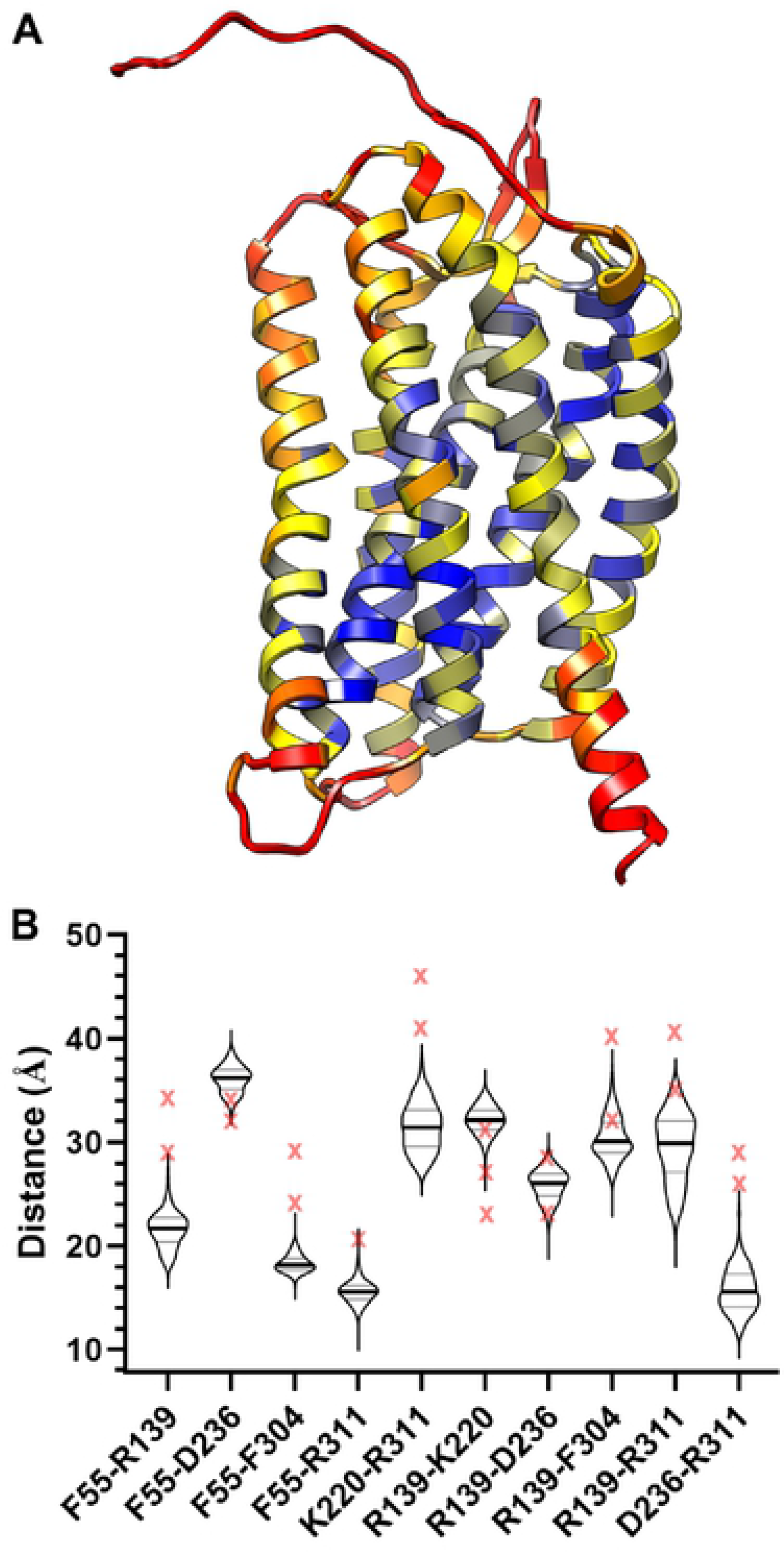
Empty AT_1_R model. **A**, The empty AT_1_R model with the movement of each residue color-coded based on the RMSF. The minimum observed RMSF (0.796 Å) is blue, 25^th^ percentile (1.689 Å) is yellow, median (2.272 Å) is orange, and the 75^th^ percentile (3.165 Å) and higher is red. **B**, A violin plot of the distances (Å) between the center of mass of the residues used in DEER spectroscopy(14) over the last 20 ns of simulation. The thick line represents the median and the thinner lines the 25^th^ and 75^th^ percentile. The red X represents the major modes from the DEER spectroscopy.

In theory, it is possible that an active empty receptor conformation could be obtained by removal of the inverse agonist olmesartan followed by MD simulation, especially since the AT_1_R is known to display basal activity. In order to confirm that the empty AT_1_R model is in an inactive conformation, consensus GPCR changes(15) between inactive and active structures were measured over the last 20 ns of each simulation (Table 1). No significant conformational change from the crystalized AT_1_Rs was observed. However, the consensus distance metrics measured from the AT_1_R simulations and the crystalized AT_1_Rs are 1.7 to 2.2 Å larger than other crystalized inactive GPCRs. Despite the difference in the distances, structural alignment of the DRY, PIF, and NPxxY motifs between the empty AT_1_R model and inactive A_2A_R crystal structure (PDB:3EML) demonstrate that the empty AT_1_R model is inactive (Figure 2). The NPxxY motif is not as clearly aligned as the DRY and PIF, but this is primarily due to the unique position of helix 8 in the empty AT_1_R model. For further comparison, the empty AT_1_R model was aligned to the active mu-opioid receptor (PDB:5C1M), demonstrating the drastic shifts in the DRY, PIF, and NPxxY required to assume the active state (Supplemental Figure 2). Therefore, the empty AT_1_R model is a model of the inactive human AT_1_R.

**Table 1:** Comparison of the empty AT1R model structure to consensus changes in GPCR structure when activated and the AT1R crystal structures. All measurements are in angstroms.

| Interaction | Inactive GPCRs* |  | Active GPCRs* |  | AT<sub>1</sub>R Crystal structures† |  | AT<sub>1</sub>R model (last 20 ns)‡ |  |
| --- | --- | --- | --- | --- | --- | --- | --- | --- |
|  | Mean | SD | Mean | SD | 4ZUD | 4YAY | Mean | SD |
| 3.46 to 6.37 | 3.8 | 0.2 | 11.0 | 2.9 | 5.5 | 5.6 | 6.0 | 0.2 |
| 1.53 to 7.53 | 3.8 | 0.1 | 9.9 | 1.4 | 6.0 | 6.1 | 5.5 | 0.3 |
| 5.55 to 6.41 | 9.2 | 1.5 | 3.8 | 0.3 | 13.1 | 12.6 | 13.3 | 0.4 |
| 3.46 to 7.53 | 9.0 | 1.1 | 3.6 | 0.2 | 11.8 | 12.7 | 12.5 | 0.2 |

\* Based on data from A.J Venkatakrishnan et al.(15) the identified residues were measured in inactive (1GZM, 2RH1, 3EML, 3UON, and 4DKL) and matching active (3PQR, 3SN6, 2YDV, 4MQS, and 5C1M) receptor PDBs; n = 5.
† There are only two inactive AT<sub>1</sub>R crystal structures, and the PDB codes are provided in the table.
‡ The distance was calculated over the last 20 ns for each rep resulting in three average values, which were then averaged; n = 3.

**Table 2.** Comparison of experimental and docking affinities to polymorphic AT_1_R.

| nsSNP<br>(Ref) | ARB used<br>experimentally | Experimental<br>Ratio* | Optimized<br>Parameters<br>Docking Ratio* | Within 2-fold the<br>error? |
| --- | --- | --- | --- | --- |
| A163T<br>(16) | Candesartan | 0.94 ± 0.09 | 21.24 ± 16.52 | No |
|  | Irbesartan | 1.84 ± 0.22 | 2.52 ± 0.14 | Yes |
|  | EXP3174 | 7.81 ± 1.12 | 3.43 ± 0.69 | No |
|  | Telmisartan | 0.70 ± 0.14 | 0.50 ± 0.15 | Yes |
|  | Valsartan | 1.10 ± 0.22 | 3.25 ± 1.21 | Yes |

\* Ratios were determined by dividing the reported data for nsSNP by wild-type, and error was calculated through error propagation using the standard deviation.

**Figure 2:**
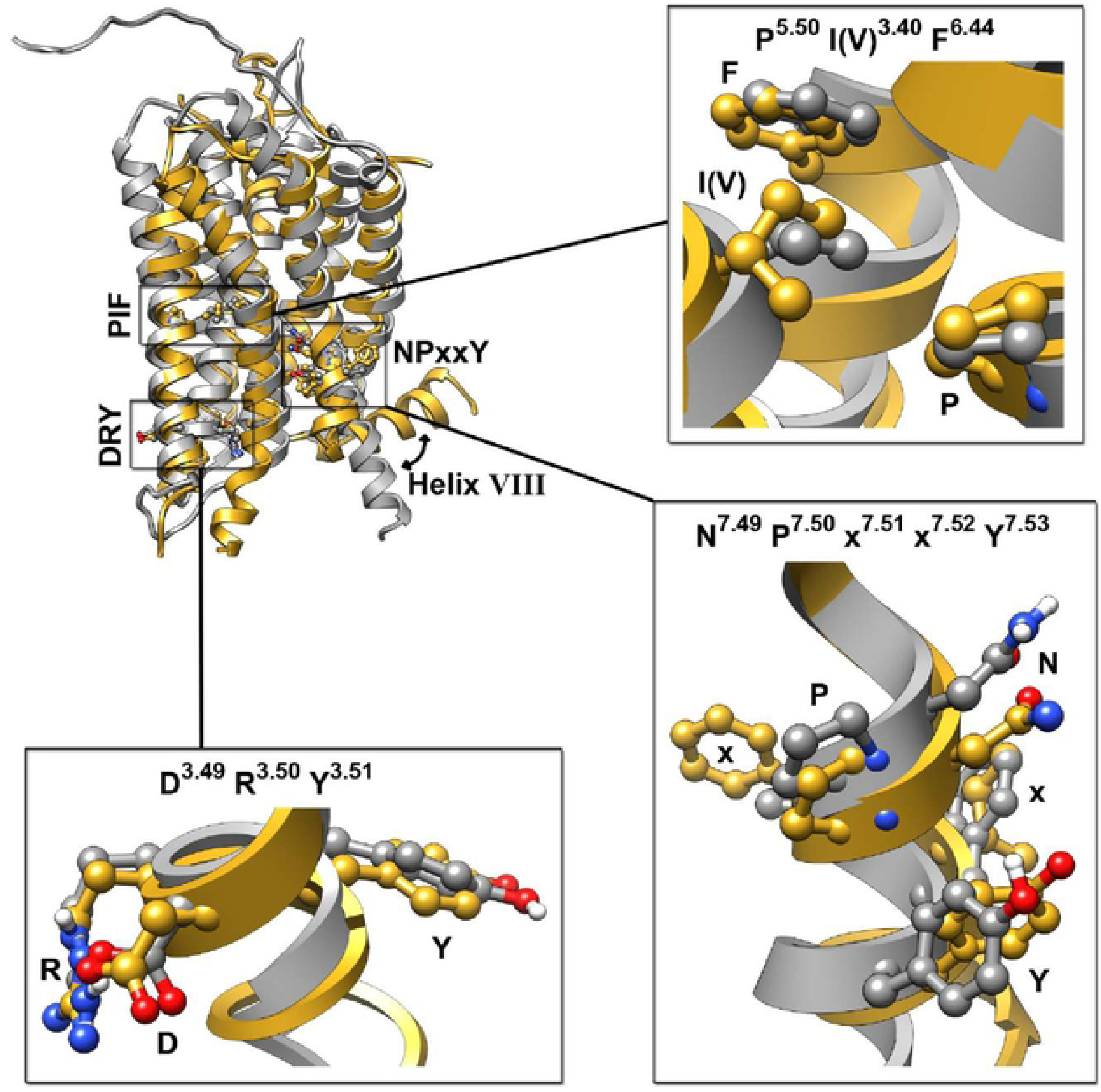
Comparison of the empty AT_1_R model to the inactive A_2A_R crystal structure (3EML). The AT_1_R (grey) is aligned to the A_2A_R (goldenrod), and the activation motifs are expanded in the boxes with Ballesteros-Weinstein numbering.

Since removing olmesartan resulted in a change in the binding pocket, the integrity of the binding pocket of the empty AT_1_R model was examined to ensure that the critical residues involved in binding are still oriented in a manner facilitating binding.(8) As shown in Figure 3a, the residues interacting with all ARBs (shown as sticks) still orientate toward the central pocket. Residues involved in some, but not all, ARB binding (shown as lines) also line the pocket. As expected due to induced fit, the binding pocket of the empty AT_1_R model is smaller than the source model 4ZUD (Figure 3B), indicating that the movement observed (Figure 1b) is the expected relaxation of the binding pocket. Fortunately, the residues involved in ARB binding remain oriented towards the binding pocket allowing for docking experiments to the empty AT_1_R model.

**Figure 3:**
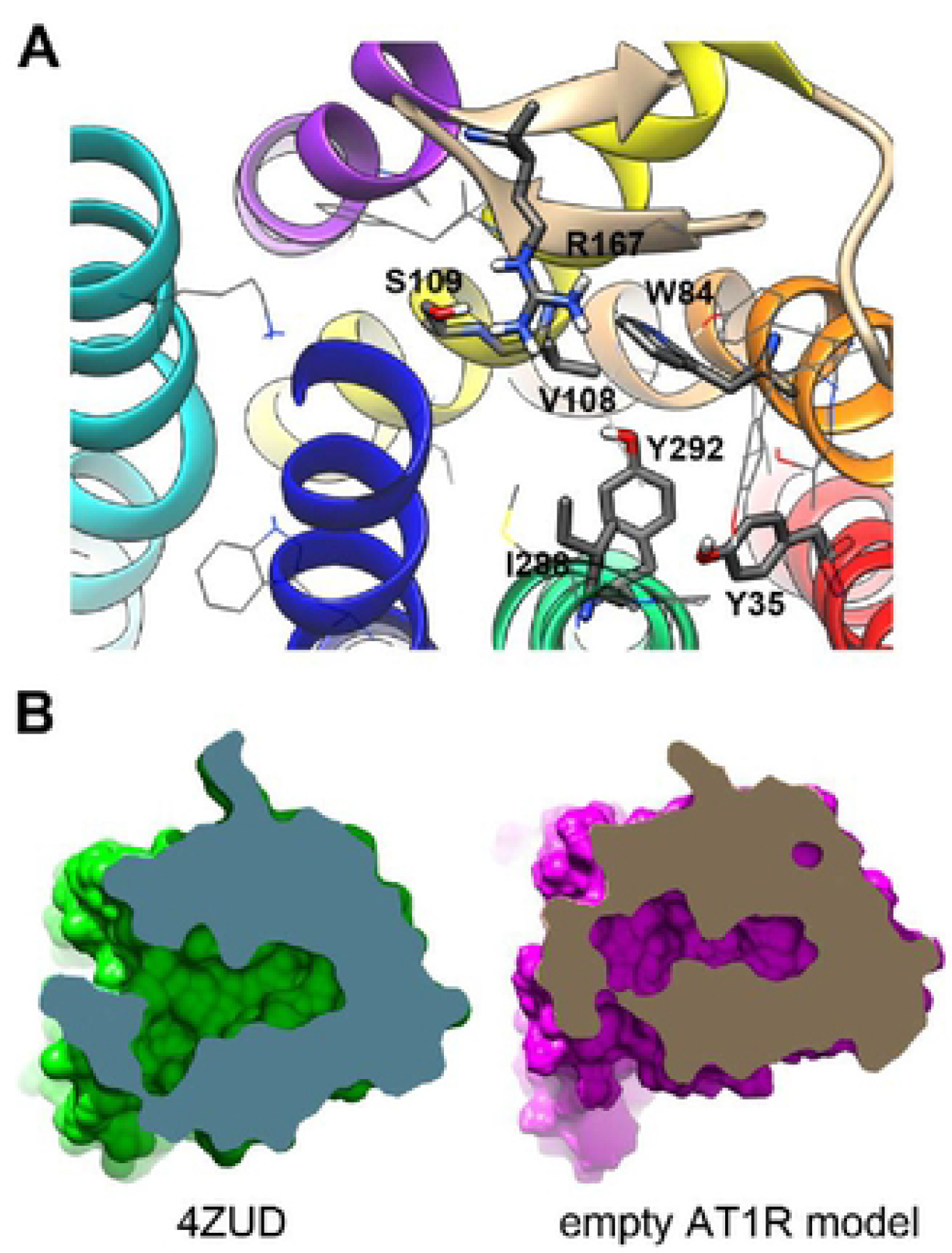
ARB binding pocket within the empty AT_1_R model. **A**, All residues shown to bind to ARBs remain directed toward the ligand-binding pocket; residues predicted to interact with all ARBs are shown as tubes and residues interacting with a subset of ARBs are shown as sticks. However, **B**, relaxation of the binding pocket during the simulation reduced its volume due to the absence of olmesartan. 4ZUD (green with a blue cutaway) and the empty AT_1_R model (purple with brown cutaway) were superimposed and slices made in UCSF Chimera.

### Docking ARBs to the AT_1_R Model

Initially, docking was conducted using standard AutoDock parameters; the search space was based on an alignment of the empty AT_1_R model with 4ZUD containing olmesartan. As shown in the blue points within Figure 4a, the weighted average affinity of three separate runs of AutoDock did not consistently generate affinities similar to known values. The AutoDock predicted affinity for four ARBs (eprosartan, irbesartan, olmesartan, and telmisartan) was below the lowest reported experimentally derived affinity, and three ARBs (azilsartan, candesartan, and EXP3174, the active metabolite of losartan) were outside of 50% of all reported affinities. Only the predicted affinity for valsartan matched experimental data.

**Figure 4:**
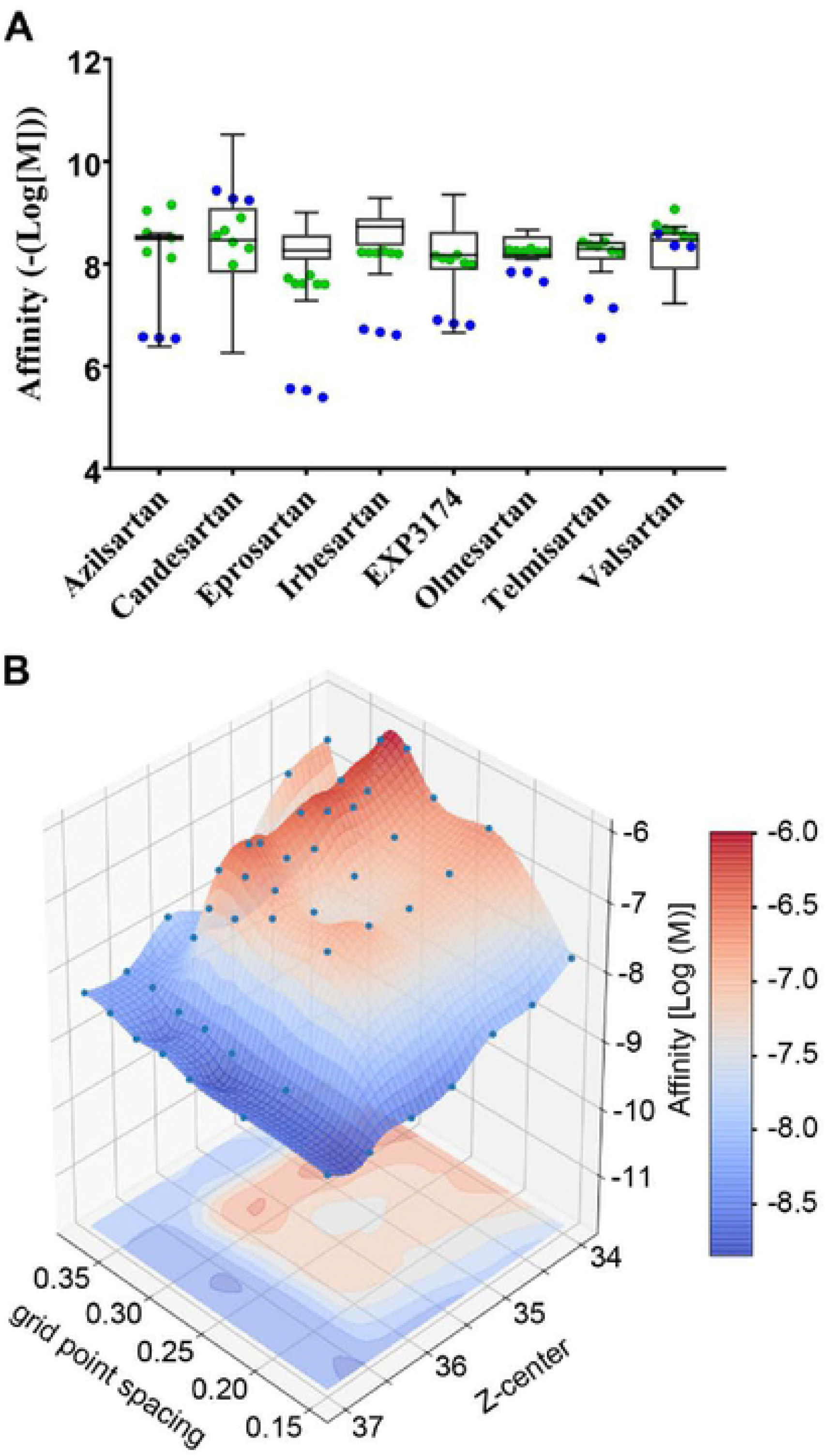
Docking ARBs to the empty AT_1_R model. **A**, Original AutoDock parameters (blue points) and optimized docking parameters (green points) are overlaid on a box and whisker plot of experimentally derived ARB affinity.(24) The whiskers represent the maximum and minimum affinity, and the box represents the second and third quartiles with the line representing the median. **B**, 3D fitting to optimize AutoDock grid point spacing and grid box center for olmesartan binding to the empty AT_1_R model.

As the predicted AutoDock affinities varied from experimental data, pilot projects were established with olmesartan to determine the AutoDock settings that could be altered to improve the predicted affinity. First, the grid spacing of the search space was systematically decreased from the default of 0.375 Å to 0.15, resulting in a non-linear relationship between grid spacing and predicted binding affinity. Second, the Z-center was altered by 0.5 Å steps in both directions resulting in a non-linear relationship between Z-center and affinity. Therefore, a seven-by-seven matrix of affinities obtained at different grid spacing and Z-axis were created for each ARB and analyzed via multiple three-dimensional fitting parameters as described in the methods. Figure 4b displays the cubic interpolation of the olmesartan seven-by-seven matrix. Each three-dimensional fit was then interrogated to identify the grid spacing and Z-axis values that generated a predicted affinity closest to the experimentally derived median affinity for each ARB. Fortunately, each three-dimensional fit produced only one set of values predicted to generate the known median affinity. The predicted values were then tested in AutoDock (n = 6), and the results that most closely mirrored the known affinities were used as the coordinates for all future docking. Supplemental Figure 3 displays the plots that generated the coordinates for each ARB.

The optimized docking parameters (Figure 4a, green points) were compared to the original docking parameters (Figure 4a, blue points). The optimized parameters more reliably predict known ARB affinity. The mean of all the optimized parameters fell within the reported range of the affinity for the respective ARB. Moreover, the mean values of the optimized docking were only separated from the median experimentally derived affinity by 2.01 ± 2.08 nM compared to 504 ± 1116 nM when using the standard settings. The highest deviation from the known median ARB affinity was eprosartan in both data sets (12.34 ± 2.12 nM in the optimized set versus 3,254 ± 711 nM in the standard set), providing an example of how the optimization protocol improved the data.

### Docking ARBs to polymorphic AT_1_Rs

After optimizing the parameters to produce affinities inline with experimentally derived affinities, the *agtr1* nsSNPs from the 1000 genome database were mapped to the empty AT_1_R model (Figure 5). The polymorphisms were introduced into the empty AT_1_R model while in a POPC/cholesterol membrane and immediately minimized within Molecular Operating Environment (MOE) software. Each empty polymorphic AT_1_R was aligned to the wild-type empty AT_1_R, and the energy minimized coordinates were utilized to conduct docking as described previously. The data summarized in Figure 6 represent affinities reduced by 2-fold or more and statistically different from the affinity for the given ARB to the wild-type empty AT_1_R. No affinities statistically increased by 2-fold or more than the control. The data predicts that many polymorphisms alter ARB affinity, but few polymorphisms adversely affect the affinity of all ARBs.

**Figure 5:**
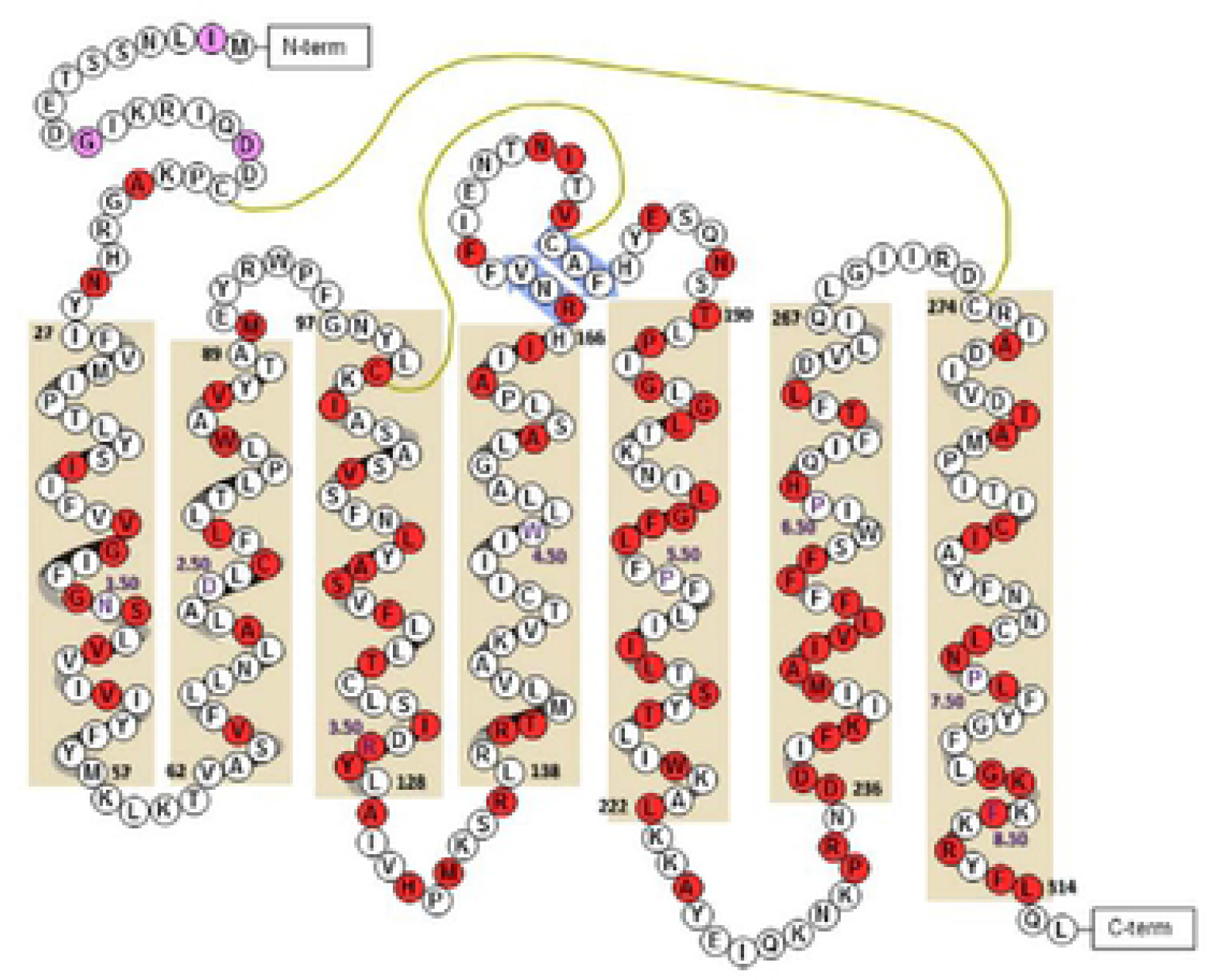
Location of hAT_1_R nsSNPs. A cartoon of the empty AT_1_R model with alpha helixes depicted within tan boxes, beta sheets depicted within blue arrows, di-sulfide bonds as mustard-colored lines, and residue numbers displaying the residue id of the beginning and end of each helix in black, as well as the Ballesteros-Weinstein numbering of the most conserved residue in purple text. Each polymorphism is shaded pink, if not studied, or red, if examined in this manuscript. The carboxyl unstructured region was not included and contains an additional 15 nsSNPs.

**Figure 6:**
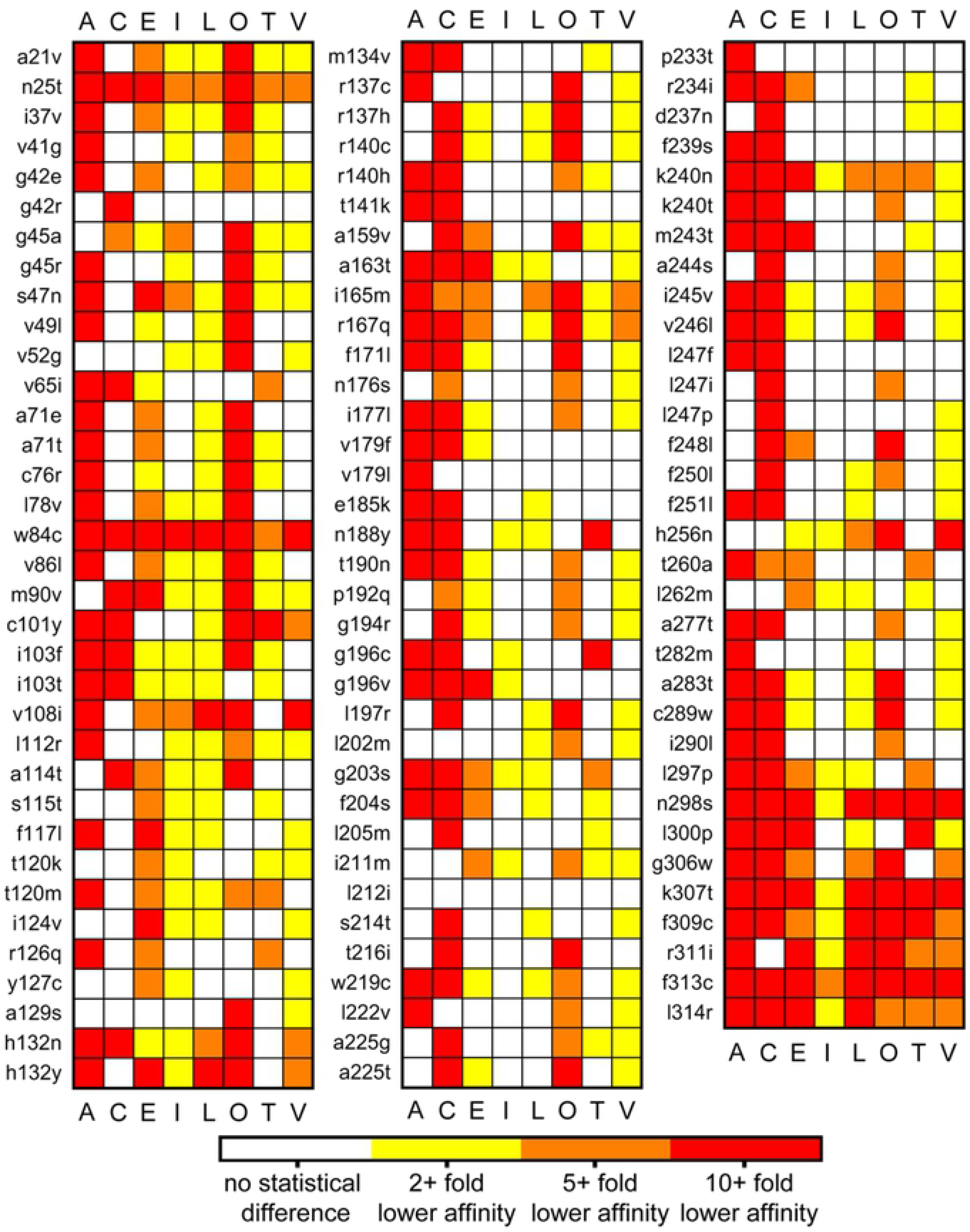
Predicted effect of AT_1_R nsSNPs on ARB affinity. Each ARB, denoted by the first letter of their name with EXP3174 marked as L for Losartan, was docked to known polymorphisms, listed on the left of each column, within the empty AT_1_R model. Data are colored a per the key as a combination of fold change and statistical difference from wild type affinities as determined by a one-way ANOVA followed by Kruskal-Wallis Multiple-Comparison Z-Value Test, n = 6.

Multiple mutagenesis studies of the AT_1_R coupled to radioligand displacement assays have been conducted since the AT_1_R was cloned; however, only a few studies utilized mutations that correspond to known nsSNPs of the AT_1_R.(9, 10, 16) Mutagenesis studies that match known polymorphisms can be compared to the predicted affinity from AutoDock to indicate the predictive power of the models (Table 3). Arsenault et al. specifically examined the Kd of ARBs to A163T hAT_1_R and found particular resistance to losartan, EXP3174, and irbesartan, as well as a higher affinity for telmisartan.(16) The docking data presented in Figure 6 and Table 3 correspond to lower affinities for EXP3174 and irbesartan as well as suggest that telmisartan binds more readily. However, the wild-type hAT_1_R:A163T hAT_1_R ratio of predicted affinities for candesartan and valsartan overestimate the change in affinity by 23 ± 18 fold and 3.0 ± 1.3 fold, respectively. Although there is little experimental data for direct comparison, the docking correctly predicted the direction of the change in affinity 60% of the time and was within two times the calculated error 60% of the time. Therefore, using MOE minimization followed by ARB docking via AutoDock resulted in a likely predictive rate of approximately 60% to identify a patient as resistant to a specific ARB.

**Table 3.** Optimal AutoDock parameters identified to obtain known median affinity for each ARBs to the empty AT_1_R model.

| ARB | Median Affinity* |  | Gridpoint<br>Spacing | X-center | Y-center | Z-center | Number of Points |  |  |
| --- | --- | --- | --- | --- | --- | --- | --- | --- | --- |
|  | Log<br>(M) | nM |  |  |  |  | X | Y | Z |
| Azilsartan | -8.51 | 3.09 | 0.168 | 38 | 61 | 36.74 | 126 | 126 | 70 |
| Candesartan | -8.46 | 3.47 | 0.342 | 38 | 61 | 34.38 | 50 | 50 | 30 |
| Eprosartan | -8.26 | 5.50 | 0.161 | 38 | 61 | 36.94 | 126 | 126 | 70 |
| Irbesartan | -8.72 | 1.91 | 0.168 | 38 | 61 | 36.84 | 126 | 126 | 70 |
| Losartan<br>(EXP3174) | -7.71 | 19.50 | 0.150 | 38 | 61 | 37.00 | 126 | 126 | 70 |
| Olmesartan | -8.17 | 6.76 | 0.357 | 38 | 61 | 35.55 | 55 | 55 | 30 |
| Telmisartan | -8.17 | 6.76 | 0.275 | 38 | 61 | 36.50 | 70 | 70 | 40 |
| Valsartan | -8.3 | 5.01 | 0.318 | 38 | 61 | 35.33 | 50 | 50 | 30 |

\* The median affinity is derived from the data presented in Figure 4A, which is from Michel MC et al.(24)

## Discussion

The empty AT_1_R model is the third ligand-free molecular dynamics derived model created after the AT_1_R was crystallized.(14, 17) In the crystalized AT_1_Rs (4ZUD and 4YAY) Arg167 points toward the ligand-binding pocket;(7, 8) however, unpublished homology modeling followed by MD simulation of the AT_1_R from our group and similarly published studies orientate Arg167 away from the binding pocket.(18, 19) Arg167 is essential for Ang II binding based on mutagenesis and modeling studies (7, 11, 17, 20) and predicted to be essential for ARB binding based on the crystal structures and subsequent docking.(8) Furthermore, this study also predicts that Arg167 is involved in the binding of most ARBs (Figure 6). Therefore, homology-based AT_1_R models that have Arg167 oriented away from the binding pocket are not accurate, and ligand binding to homology models with Arg167 in the wrong orientation are subsequently inaccurate. Based on the orientation of crucial ARB binding residues within the model empty AT_1_R (Figure 3A), the ligand-binding pocket of the empty AT_1_R model is likely an accurate model of the ligand-free AT_1_R.

Although the empty AT_1_R model is in the inactive state (Table 1 and Figure 2), the flexibility of helix 8 was surprising. Helix 8 is not present in 4ZUD,(7) but is in 4YAY(8) as well as the active AT_1_R structure 6DO1.(20) The AT_1_R crystal structures place helix 8 slightly bent from the membrane and parallel to the membrane, respectively. Fortunately, the mobility of helix 8 is similar to what was measured via DEER spectroscopy and MD modeling.(14, 17) Helix 8 is flexible without negatively charged phospholipids;(14) however, the flexibility of helix 8 is likely restrained in vivo due to the C-terminus anchoring to negative phospholipids in the membrane.(17, 21–23) The empty AT_1_R system presented here lacks lipids shown to enhance helix 8 association with the membrane as the modeled membrane only contains POPC and cholesterol.(21–23) Additionally, 4YAY, which served as a guide, does not contain the entirety of helix 8; thus, the empty AT_1_R model lacks four positively charged amino acids that facilitate binding to negative phospholipids.(23) Therefore, the mobility of helix 8 in the empty AT_1_R model may be dependent on its length and the lack of negative phospholipids. Future MD simulations of the AT_1_R, and likely other Gq-coupled GPCRs, should be conducted in a membrane with PIP2 as PIP2 is the substrate of PLC and is expected near Gq-coupled GPCRs.

Given that the ligand-binding site of the empty AT_1_R model appears appropriate, large scale docking began to assess many of the known *agtr1* nsSNPs (Figure 5). However, before testing all the nsSNPs, the docking had to be optimized to produce orientations of the ARB that resembled the crystal structure, specifically for olmesartan,(7) and affinities matching known affinity for each ARB.(24) AutoDock was chosen specifically for the ability to obtain an estimated affinity, and as shown in Figure 4, the original parameters did not produce affinities within the known range for olmesartan although the orientation was accurate (not shown). Therefore, the parameters were optimized through non-linear topographical methods resulting in an improved affinity prediction for all ARBs. There is a danger in the method described, as the highest affinity predicted was often greater than known median affinity and occasionally outside the reported range of affinities for a specific ARB. Therefore, this method should only be used to tune AutoDock settings to obtain known affinities.

### Role of molecular modeling in personalized medicine

Achievable personalized medicine can occur through matching the pharmacopeia to a person’s genome. For example, polymorphisms within metabolic enzymes are accepted measures to adjust the dosing of a drug or inform clinicians to use an alternative medication to avoid a potential drug-gene interaction. The next steps to bring personalized medicine to the clinic include understanding how polymorphisms alter individual drug effects. Polymorphisms can alter the affinity of a drug as well as alter signal transduction, as exemplified by the example of V302I μ-opioid receptors that results in the ablation of naloxone function.(6) Herein, the data suggest that polymorphisms in the AT_1_R may be responsible for apparent ARB resistance. Moreover, the data suggests that often there is an ideal ARB for a particular polymorphism; thus, matching a patient’s genotype to an existing drug fulfills the promise of personalized medicine.

Using computational modeling provides a relatively inexpensive and rapid drug screening process. As technology progresses, the speed of the computations increases; whereas, fundamental pharmacological and biochemical determination of ligand affinity has changed little, and radio-labeled ligands are not always available. Importantly, the nsSNPs of the AT_1_R examined are only the tip of the proverbial iceberg. There is little data regarding the combination of nsSNPs, and with 103 polymorphisms, the combinatorial potential is staggering. Moreover, Hauser et al. reported that a fraction over 1 in 293 births contain a new nsSNP in a clinically targeted GPCR.(6) Currently, there are an estimated 250 births worldwide per minute; thus, nearly every minute there is a new polymorphism created in one of the 108 GPCRs that are clinical targets. Therefore, the number of seemingly benign nsSNPs in the AT_1_R that may attenuate ARB affinity will likely increase in the future. A computational approach to ARB affinity is a reasonable and comparatively rapid method to tailor ARB therapy to a patient. Ideally, a patient’s *agrt1* would be sequenced and compared to a database to select the best ARB. Any novel nsSNPs, or nsSNP combinations, would then be modeled to predict which ARB is most appropriate and after verification added to the database.

### Limitations of the Current Study

Although the affinity estimates for each ARB blinding to the empty AT_1_R model are within known ranges and comparison of relative affinities obtained through molecular modeling is standard practice, there are caveats in the experimental design that prevents taking the data directly to the clinic. The caveats include a lack of AT_1_R flexibility in the docking algorithm; lack of data on Ang II affinity to compare to ARB affinity, as ARB efficacy is dependent on the ratio of the ARB to Ang II affinity; and lack of a biological understanding of each polymorphism. AutoDock allows for limited flexible residues, but do not account for the flexibility of the entire receptor,(25) and Ang II has too many rotatable bonds to dock reliably in AutoDock. MD simulation allows for all-atom flexibility and estimation of ARB and Ang II affinity, but the computational time required to examine each ligand and polymorphism is beyond the scope of a single laboratory. Additionally, polymorphisms can have a myriad of effects such as altering the surface expression or blocking coupling to the G-protein. Biological experiments are necessary to establish the functional relevance of each polymorphism as some polymorphisms may ablate the function of the AT_1_R. Therefore, to realize personalized drug therapy, modeling and basic laboratories can collaborate to create viable databases to guide clinical decisions.

## Methods

### AT_1_R Model preparation

The crystal structure of human AT_1_R bound to olmesartan (PDB: 4ZUD) was downloaded from the RCSB Protein Data Bank.(7) 4ZUD contains apocytochrome b_562_RIL fused to the amino terminus, and many of the flexible regions, as well as helix 8, of the AT_1_R are not resolved. In order to generate an appropriate starting structure, olmesartan and the apocytochrome b_562_RIL fusion were removed from 4ZUD, and the missing regions were added to the protein with MOE software (Chemical Computing Group ULC, Montreal, Canada). Specifically, the N-Terminus (residues 1 to 25), intracellular loop 2 (residues 134 to 140), extracellular loop 2 (residues 186 to 188), intracellular loop 3 (residues 223 to 234), and helix 8 (residues 305 to 316) were added to the AT_1_R in accordance to the human AT_1_R sequence and PDB:4YAY. The remaining carboxyl-tail of the AT_1_R (residues 317 to 359) was not modeled. The AT_1_R model then underwent an energy minimization within MOE using the Amber10:Extended Huckel Theory (EHT) force field.(26)

### Molecular dynamic (MD) simulations

The MOE minimized AT_1_R was loaded into CHARMM-GUI.(27) An 80 Å by 80 Å lipid bi-layer composed of 13% cholesterol(28) and 87% Phosphatidylcholine (POPC) was generated around the receptor. Water was packed 17.5 Å above and below the lipid bi-layer, and 150 mM Na^+^ and Cl^−^ ions were added to the system via Monte-Carlo ion placing. The all-atom CHARMM C36 force field(29) for proteins and ions, and the CHARMM TIP3P force field(30) for water were selected. A hard non-bonded cutoff of 8.0 angstroms was utilized. All molecular dynamics simulations were performed using the PMEMD module of the AMBER16 package(31) with support for MPI multi-process control and GPU acceleration code. Orthorhombic periodic boundary conditions with a constant pressure of 1 atm was set via the NPT ensemble and temperature was set to 310.15°K (37°C) using Langevin dynamics. The SHAKE algorithm was used to constrain bonds containing hydrogens. The dynamics were propagated using Langevin dynamics with Langevin damping coefficient of 1 ps^−1^ and a time step of 2 fs. Before the production run, the AT_1_R model was minimized for 5000 steps using the steepest descent method and then equilibrated for 600 ps. The protein coordinates were saved in 10 ps intervals. The production run lasted 150 ns, at which point all three replicas were stable for at least 20 ns. The frame representing the value closest to the average RMSD of the stable 20 ns from replica 1 was selected as the structure representing the empty conformation of the AT_1_R and herein is called the empty AT_1_R model.

Since the AT_1_R has been extensively studied and is known to display basal activity in the absence of agonist,(32) the empty AT_1_R model was examined to determine if it resembled an inactive or active GPCR. Venkatakrishnam et al. identified residues that are commonly in contact when a GPCR is inactive but not active and vice versa;(15) based on the identified interactions within the crystal structures, the atoms nearest each other were identified and the distances measured in PyMOL software in the inactive and active state. The same measurements were made in the two AT_1_R crystal structures (4ZUD and 4YAY) and over the last 20 ns of each of the three replica MD simulations. Wingler et al. conducted DEER spectroscopy of apo-AT_1_R,(14) similar measurements were made via measuring the distance between the center of mass of the residues examined in the DEER spectroscopy studies across the entire simulation utilizing CPPTRAJ.(33)

### Docking of ARBs to the empty AT_1_R

The empty AT_1_R and all eight clinically viable ARBs, as mol2 files, were loaded individually into AutoDock Tools 1.5.6 and docking was conducted as described by S. Forli et al.(25) The metabolite of losartan, EXP3174, is a potent inhibitor of the AT_1_R;(34) therefore, EXP3174 was utilized instead of losartan. In brief, Gasteiger charges were computed, and all atoms were assigned an AD4 type. Ligand torsions were calculated, and all non-polar hydrogens on the protein were merged before assigning atom types. When preparing for docking, the ligand-binding pocket was selected to encompass the olmesartan binding pocket from 4ZUD. Each ligand was docked to the empty AT_1_R model 100 times in a single AutoDock 4.0 run, and each run was replicated three times. A Boltzmann weighted average affinity was calculated for each run and converted from docking estimated binding free energy (*ΔG*) in kcal/mol to a predicted dissociation constant (*K*_*d*_) via Equation 1, shown below. Note (*i*) is the index for any given affinity from AutoDock, and RT is the molar gas constant and temperature in kelvin, respectively.

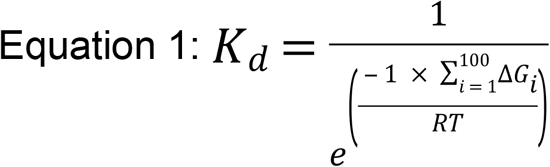

Although the optimal binding site matched previous data,(8) the predicted K_d_ did not match known binding affinities. Therefore, we optimized the docking parameters by performing different types of least squares curve fitting of a seven-by-seven matrix of Boltzmann weighted average binding affinities for each ARB. The variables altered were the grid box spacing (Å) (0.150, 0.225, 0.250, 0.275, 0.300, 0.325, 0.350, 0.375) and the center position of the grid box on the Z-axis, altered by 0.5 Å in the positive and negative direction by three steps from 35.5 (34.0, 34.5, 35.0, 35.5, 36.0, 36.5, 37.0). Each point in the matrix was created by a single 100 dockings run of AutoDock 4.0, and the 100 affinities were converted to a predicted K_d_ as described above. The data were then fit by a nearest, linear, cubic, and bivariate spline 3D-fitting using python (see supplemental methods for the python script). The variables resulting in the greatest predicted K_d_, as well as the known median ARB affinity,(24) were extracted from the curve fitting and tested in Autodock 4.0 three times to confirm the predicted values. For telmisartan, the seven-by-seven matrix identified ideal parameters; thus, those parameters were utilized. The grid box spacing and Z-axis that produced a predicted K_d_ closest to the experimentally derived median affinity for each ARB was run six times in Autodock; only data from the most accurate method is reported. See Table 3 for the parameters that produced a predicted K_d_ similar to the known median affinity for each ARB.

### Generating nsSNPs AT_1_R and docking ARBs

Data from the 1000s genome project was mined to extract all non-synonymous single point mutations (nsSNPs).(13) The empty AT_1_R within the lipid bilayer was loaded into MOE, and each of the 103 chosen nsSNPs was generated individually using the protein builder function in MOE software. After changing the residue, an energy minimization utilizing Amber10 EHT force field was conducted via MOE to generate an appropriate conformation of the AT_1_R carrying the polymorphism. The atom coordinates of empty polymorphic AT_1_R was saved as a PDB and aligned to the wild-type empty AT_1_R so that all docking coordinates would be similar. Each empty polymorphic AT_1_R was loaded into AutoDockTools as described previously, and the ARB specific optimized parameters were utilized to determine the affinity of each ARB to each polymorphic AT_1_R. Each ligand was docked to the receptor 100 times in a single AutoDock 4.0 run, and each run was replicated six times.

### Statistical Analysis

The fold change in binding affinity between wild-type AT_1_R and each polymorphic AT_1_R was analyzed within NCSS 2007 statistical software (Kaysville, UT) using a one-way ANOVA followed by Kruskal-Wallis Multiple-Comparison Z-Value Test. Data are shown if the fold change is 2-fold or higher and statistically different than control. No ARBs displayed a reduction by 2-fold or higher and were statistically different from control.

## Acknowledgments

The authors greatly appreciate Martin C. Michel for providing the data reported in their comprehensive review of ARBs,(24) allowing for the creation of Figure 2.

## Data Supplements

**Supplemental Figure 1: Analysis of structural stability and helix 8 movement. A**, RMSD of each model indicates that the empty AT_1_R models are stable; **B**, however, helix 8 is mobile when global movement is minimal. The empty AT_1_R model served as the reference for helix 8 mobility, and it is the only frame that orientates helix 8 as an extension of helix 7 (0 RMSD in replica 1 red line R1:h8).

**Supplemental Figure 2: Comparison of the empty AT_1_R model to the active mu-opioid receptor (5C1M).** The AT_1_R (grey) is aligned to the μOR (goldenrod), and the activation motifs are expanded in the boxes with Ballesteros-Weinstein numbering.

**Supplanted Figure 3: ARB binding optimization.** 3D fitting to optimize AutoDock grid point spacing and grid box center for ARB binding to the empty AT_1_R model. The fit providing the optimal parameters is listed below the ARB. The seven-by-seven table best-modeled telmisartan; thus, telmisartan is not shown.

**Supplemental Methods: Scripts utilized to optimize AutoDock parameters.** The script is written in MS Word and cannot be cut and pasted to run due to different handling of fonts. Notes to help the user are in red boxes on the left-hand side of the text.

**Supplemental file**: empty AT_1_R model

## References

1. Vinson GP. Why isn’t the angiotensin type 1 receptor a target in cancer? Oncotarget. 2017;8(12):18618–9.

2. Dasgupta C, Zhang L. Angiotensin II receptors and drug discovery in cardiovascular disease. Drug Discov Today. 2011;16(1-2):22–34.

3. Liu DX, Zhang YQ, Hu B, Zhang J, Zhao Q. Association of AT1R polymorphism with hypertension risk: An update meta-analysis based on 28,952 subjects. J Renin Angiotensin Aldosterone Syst. 2015;16(4):898–909.

4. Feng X, Zheng BS, Shi JJ, Qian J, He W, Zhou HF. A systematic review and meta-analysis of the association between angiotensin II type 1 receptor A1166C gene polymorphism and myocardial infarction susceptibility. J Renin Angiotensin Aldosterone Syst. 2014;15(3):307–15.

5. Spiering W, Kroon AA, Fuss-Lejeune MJ, de Leeuw PW. Genetic contribution to the acute effects of angiotensin II type 1 receptor blockade. J Hypertens. 2005;23(4):753–8.

6. Hauser AS, Chavali S, Masuho I, Jahn LJ, Martemyanov KA, Gloriam DE, et al. Pharmacogenomics of GPCR Drug Targets. Cell. 2018;172(1-2):41–54 e19.

7. Zhang H, Unal H, Desnoyer R, Han GW, Patel N, Katritch V, et al. Structural Basis for Ligand Recognition and Functional Selectivity at Angiotensin Receptor. The Journal of biological chemistry. 2015;290(49):29127–39.

8. Zhang H, Unal H, Gati C, Han GW, Liu W, Zatsepin NA, et al. Structure of the Angiotensin receptor revealed by serial femtosecond crystallography. Cell. 2015;161(4):833–44.

9. Ji H, Leung M, Zhang Y, Catt KJ, Sandberg K. Differential structural requirements for specific binding of nonpeptide and peptide antagonists to the AT1 angiotensin receptor. Identification of amino acid residues that determine binding of the antihypertensive drug losartan. Journal of Biological Chemistry. 1994;269(24):16533–6.

10. Ji H, Zheng W, Zhang Y, Catt KJ, Sandberg K. Genetic transfer of a nonpeptide antagonist binding site to a previously unresponsive angiotensin receptor. Proc Natl Acad Sci U S A. 1995;92(20):9240–4.

11. Yamano Y, Ohyama K, Kikyo M, Sano T, Nakagomi Y, Inoue Y, et al. Mutagenesis and the molecular modeling of the rat angiotensin II receptor (AT1). Journal of Biological Chemistry. 1995;270(23):14024–30.

12. Noda K, Saad Y, Kinoshita A, Boyle TP, Graham RM, Husain A, et al. Tetrazole and carboxylate groups of angiotensin receptor antagonists bind to the same subsite by different mechanisms. Journal of Biological Chemistry. 1995;270(5):2284–9.

13. Genomes Project C, Auton A, Brooks LD, Durbin RM, Garrison EP, Kang HM, et al. A global reference for human genetic variation. Nature. 2015;526(7571):68–74.

14. Wingler LM, Elgeti M, Hilger D, Latorraca NR, Lerch MT, Staus DP, et al. Angiotensin Analogs with Divergent Bias Stabilize Distinct Receptor Conformations. Cell. 2019;176(3):468–78 e11.

15. Venkatakrishnan AJ, Deupi X, Lebon G, Heydenreich FM, Flock T, Miljus T, et al. Diverse activation pathways in class A GPCRs converge near the G-protein-coupling region. Nature. 2016;536(7617):484–7.

16. Arsenault J, Lehoux J, Lanthier L, Cabana J, Guillemette G, Lavigne P, et al. A single-nucleotide polymorphism of alanine to threonine at position 163 of the human angiotensin II type 1 receptor impairs Losartan affinity. Pharmacogenetics and genomics. 2010;20(6):377–88.

17. Modestia SM, Malta de Sa M, Auger E, Trossini GHG, Krieger JE, Rangel-Yagui CO. Biased Agonist TRV027 Determinants in AT1R by Molecular Dynamics Simulations. J Chem Inf Model. 2019.

18. Clement M, Martin SS, Beaulieu ME, Chamberland C, Lavigne P, Leduc R, et al. Determining the environment of the ligand binding pocket of the human angiotensin II type I (hAT1) receptor using the methionine proximity assay. The Journal of biological chemistry. 2005;280(29):27121–9.

19. Baleanu-Gogonea C, Karnik S. Model of the whole rat AT1 receptor and the ligand-binding site. J Mol Model. 2006;12(3):325–37.

20. Wingler LM, McMahon C, Staus DP, Lefkowitz RJ, Kruse AC. Distinctive Activation Mechanism for Angiotensin Receptor Revealed by a Synthetic Nanobody. Cell. 2019;176(3):479–90 e12.

21. Hirst DJ, Lee TH, Pattenden LK, Thomas WG, Aguilar MI. Helix 8 of the angiotensin-II type 1A receptor interacts with phosphatidylinositol phosphates and modulates membrane insertion. Sci Rep. 2015;5:9972.

22. Mozsolits H, Unabia S, Ahmad A, Morton CJ, Thomas WG, Aguilar MI. Electrostatic and hydrophobic forces tether the proximal region of the angiotensin II receptor (AT1A) carboxyl terminus to anionic lipids. Biochemistry. 2002;41(24):7830–40.

23. Kamimori H, Unabia S, Thomas WG, Aguilar MI. Evaluation of the membrane-binding properties of the proximal region of the angiotensin II receptor (AT1A) carboxyl terminus by surface plasmon resonance. Anal Sci. 2005;21(2):171–4.

24. Michel MC, Foster C, Brunner HR, Liu L. A systematic comparison of the properties of clinically used angiotensin II type 1 receptor antagonists. Pharmacological reviews. 2013;65(2):809–48.

25. Forli S, Huey R, Pique ME, Sanner MF, Goodsell DS, Olson AJ. Computational protein-ligand docking and virtual drug screening with the AutoDock suite. Nat Protoc. 2016;11(5):905–19.

26. Gerber PR, Muller K. MAB, a generally applicable molecular force field for structure modelling in medicinal chemistry. J Comput Aided Mol Des. 1995;9(3):251–68.

27. Jo S, Kim T, Iyer VG, Im W. CHARMM-GUI: a web-based graphical user interface for CHARMM. J Comput Chem. 2008;29(11):1859–65.

28. Pankov R, Markovska T, Antonov P, Ivanova L, Momchilova A. The plasma membrane lipid composition affects fusion between cells and model membranes. Chem Biol Interact. 2006;164(3):167–73.

29. Best RB, Zhu X, Shim J, Lopes PE, Mittal J, Feig M, et al. Optimization of the additive CHARMM all-atom protein force field targeting improved sampling of the backbone phi, psi and side-chain chi(1) and chi(2) dihedral angles. J Chem Theory Comput. 2012;8(9):3257–73.

30. Jorgensen WL, Chandrasekhar J, Madura JD, Impey RW, Klein MLK. Comparison of simple potential functions for simulating liquid water J Chem Phys. 1983;79(2):926–35.

31. D.A. Case RMB, D.S. Cerutti, T.E. Cheatham, III, T.A. Darden, R.E. Duke, T.J. Giese, H. Gohlke, A.W. Goetz, N. Homeyer, S. Izadi, P. Janowski, J. Kaus, A. Kovalenko, T.S. Lee, S. LeGrand, P. Li, C. Lin, T. Luchko, R. Luo, B. Madej, D. Mermelstein, K.M. Merz, G. Monard, H. Nguyen, H.T. Nguyen, I. Omelyan, A. Onufriev, D.R. Roe, A. Roitberg, C. Sagui, C.L. Simmerling, W.M. Botello-Smith, J. Swails, R.C. Walker, J. Wang, R.M. Wolf, X. Wu, L. Xiao and P.A. Kollman AMBER 16. San Francisco: University of California; 2016.

32. Martin SS, Holleran BJ, Escher E, Guillemette G, Leduc R. Activation of the angiotensin II type 1 receptor leads to movement of the sixth transmembrane domain: analysis by the substituted cysteine accessibility method. Molecular pharmacology. 2007;72(1):182–90.

33. Roe DR, Cheatham TE. PTRAJ and CPPTRAJ: Software for Processing and Analysis of Molecular Dynamics Trajectory Data. Journal of Chemical Theory and Computation. 2013;9(7):3084–95.

34. Wong PC, Price WA, Jr., Chiu AT, Duncia JV, Carini DJ, Wexler RR, et al. Nonpeptide angiotensin II receptor antagonists. XI. Pharmacology of EXP3174: an active metabolite of DuP 753, an orally active antihypertensive agent. J Pharmacol Exp Ther. 1990;255(1):211–7.

